# Acute alcohol administration dampens threat-related activation in the central extended amygdala

**DOI:** 10.1101/283358

**Authors:** Juyoen Hur, Claire M. Kaplan, Jason F. Smith, Daniel E. Bradford, Andrew S. Fox, John J. Curtin, Alexander J. Shackman

**Affiliations:** Department of Psychology, University of Maryland, College Park, MD 20742 USA; Neuroscience and Cognitive Science Program, University of Maryland, College Park, MD 20742 USA; Maryland Neuroimaging Center, University of Maryland, College Park, MD 20742 USA; Department of Psychology, University of Wisconsin—Madison, 1202 West Johnson Street, Madison, WI 53706 USA; Department of Psychology, University of California, Davis, CA 9S616 USA; California National Primate Research Center, University of California, Davis, CA 9S616 USA

**Keywords:** affective neuroscience, anxiolytic, bed nucleus of the stria terminalis (BST/BNST), central extended amygdala (EAc), ethyl alcohol, fear and anxiety, functional MRI (fMRI)

## Abstract

Alcohol abuse is common, imposes a staggering burden on public health, and is challenging to treat, underscoring the need to develop a deeper understanding of the underlying neurobiology. When administered acutely, ethyl alcohol reduces threat reactivity in humans and other animals, and there is growing evidence that threat-dampening and related negative reinforcement mechanisms support the etiology and recurrence of alcohol and other kinds of substance misuse. Converging lines of evidence motivate the hypothesis that these effects are mediated by the central extended amygdala (EAc)—including the central nucleus of the amygdala (Ce) and bed nucleus of the stria terminalis (BST)—but the relevance of this circuitry to acute alcohol effects in humans remains poorly understood. Using a single-blind, randomized-groups design, multiband imaging data were acquired from 49 social drinkers while they performed an fMRI-optimized emotional-faces/places paradigm after consuming alcohol or placebo. Relative to placebo, alcohol significantly dampened reactivity to threat-related emotional faces in the BST. To rigorously assess potential regional differences in activation, data were extracted from anatomically defined Ce and BST regions-of-interest. Analyses revealed a similar pattern of dampening across the two regions. In short, alcohol acutely dampens reactivity to threat-related faces in humans and it does so similarly across the two major divisions of the EAc. These observations provide a framework for understanding the translational relevance of addiction models derived from work in rodents, inform on-going debates about the functional organization of the EAc, and set the stage for bi-directional translational models aimed at developing improved treatment strategies for alcohol abuse and other addictions.

## INTRODUCTION

Alcohol abuse is common (i.e., nearly three-quarters of Americans consumed some form of ethanol in the past year and, among them, 17.5% met criteria for an alcohol use disorder); contributes to a wide range of adverse social outcomes (e.g., crime); and imposes a substantial and growing burden on global public health and the economy ^1–3^. Existing treatments are incompletely effective ^4^, underscoring the urgency of developing a clearer understanding of the neural systems contributing to the development, maintenance, and recurrence of alcohol abuse ^5^. When administered acutely, alcohol has anxiolytic properties in humans and rodents ^6–8^, and there is clear evidence that threat- and stress-dampening effects (i.e., negative reinforcement) contribute to the etiology and recurrence of alcohol misuse and abuse ^8, 9^. Yet, remarkably little is known about the neural circuitry underlying the threat-dampening effects of acute alcohol administration in humans.

Converging lines of mechanistic, anatomical, physiological, and pharmacological evidence highlight the potential importance of the central extended amygdala (EAc), including the central nucleus of the amygdala (Ce) and bed nucleus of the stria terminalis (BST). The EAc plays a critical role in assembling defensive responses to a range of threats ^10–13^. Anatomically, the EAc is poised to govern vigilance and other aspects of fear and anxiety via dense mono- and poly-synaptic projections to downstream effector regions ^14,15^. Imaging and lesion studies demonstrate that the EAc plays a key role in selecting and prioritizing the processing of ‘threat-related’ cues, such as fearful faces ^16^ (see **Supplementary Comment**). EAc function co-varies with individual differences in anxious temperament and likely contributes to the development and maintenance of anxiety disorders ^12,15–17^. Conversely, acute administration of classic anxiolytics (e.g., diazepam, lorazepam) is associated with reduced reactivity of the dorsocaudal amygdala (in the region of the Ce) to threat-related faces ^18,19^.

Despite this progress, the relevance of the EAc to alcohol-induced threat-dampening in humans remains poorly understood. Although a few functional MRI (fMRI) studies have been reported (Table 1), with some suggesting that alcohol dampens amygdala reactivity to threat-related faces (i.e., fearful or angry expressions), they are limited by small samples (*N*s<15) and coarse spatial resolution. Many relied on thresholding procedures that are now known to markedly inflate the risk of false discoveries ^20,21^. Often, null results were interpreted as evidence of threat-dampening ^22^. None explicitly examined either the Ce or the BST ^23–26^. All relied on randomized cross-over designs, despite evidence questioning the retest reliability of amygdala activation e.g., ^27^. In short, it remains unclear whether the Ce and BST are dampened by alcohol and, if so, whether they differ in their sensitivity.

**Table 1.**
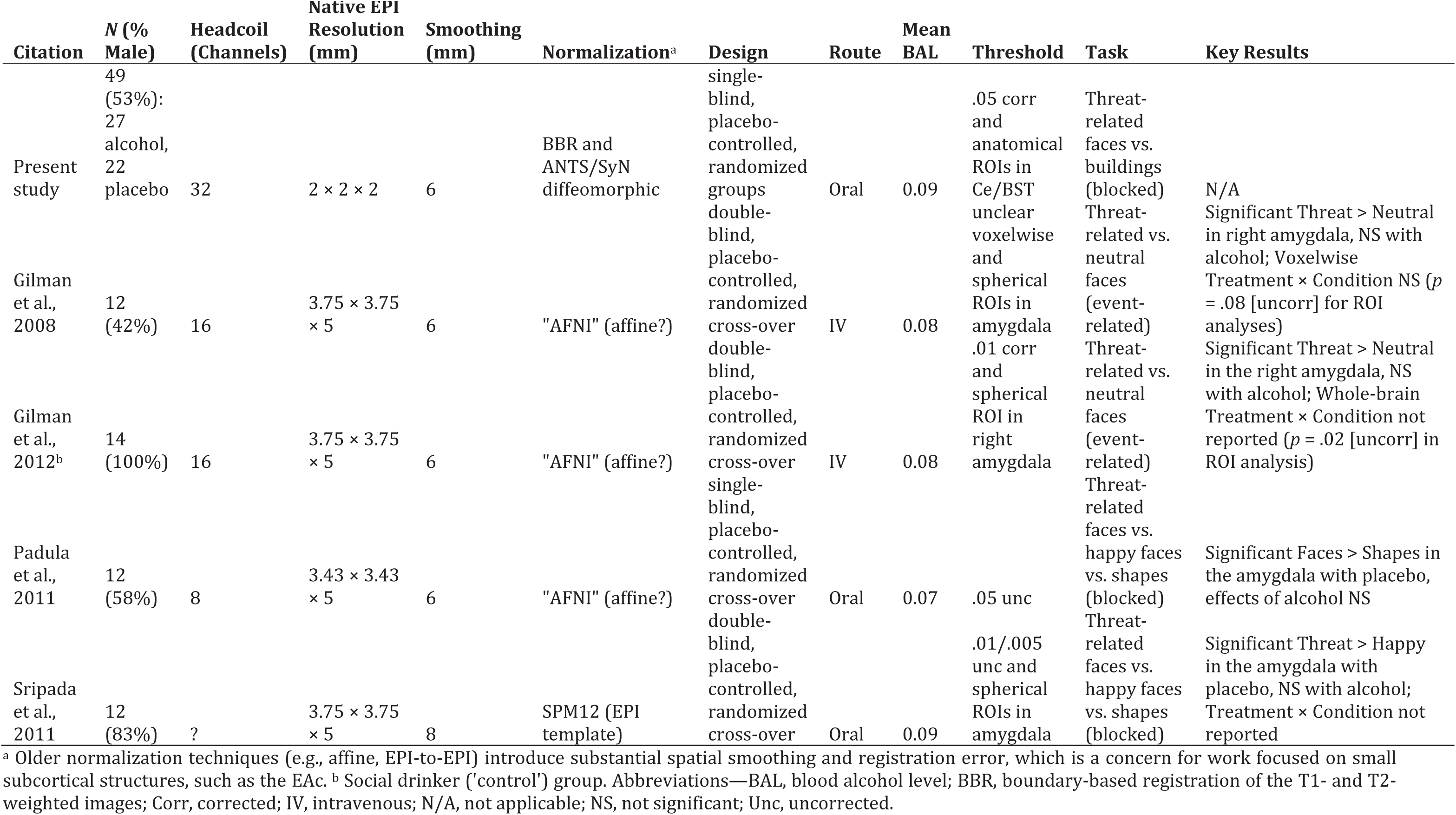
The effects of acute alcohol administration on reactivity to threat-related faces in human imaging studies.

Here, we used a novel combination of approaches to assess the influence of acute alcohol administration on reactivity to threat-related faces in the Ce and the BST. Using a single-blind, placebo-controlled, randomized-groups design (Table 2), fMRI data were acquired from 49 psychiatrically healthy social drinkers while they performed an fMRI-optimized emotional-faces/places paradigm after consuming an alcoholic or placebo beverage. Several methods enhanced precision, including a multiband pulse sequence and advanced co-registration and spatial normalization techniques ^28^. Recently developed, anatomically defined Ce and BST regions-of-interest (ROIs) ^29,30^ made it possible to directly compare the hypothesized threat-dampening effects of alcohol in the BST and the Ce for the first time. Understanding the role of the EAc in alcohol-induced threat-dampening is important. It is a necessary step to determining the translational relevance of addiction models derived from rodent models e.g., ^31^. It would also provide insight into the EAc’s role in social drinking and other kinds of substance use, inform on-going debates about the functional organization of the EAc ^12,32,33^, and guide the development of bi-directional translational models ^12,34^ aimed at developed improved treatment strategies ^5,35^.

**Table 2.**
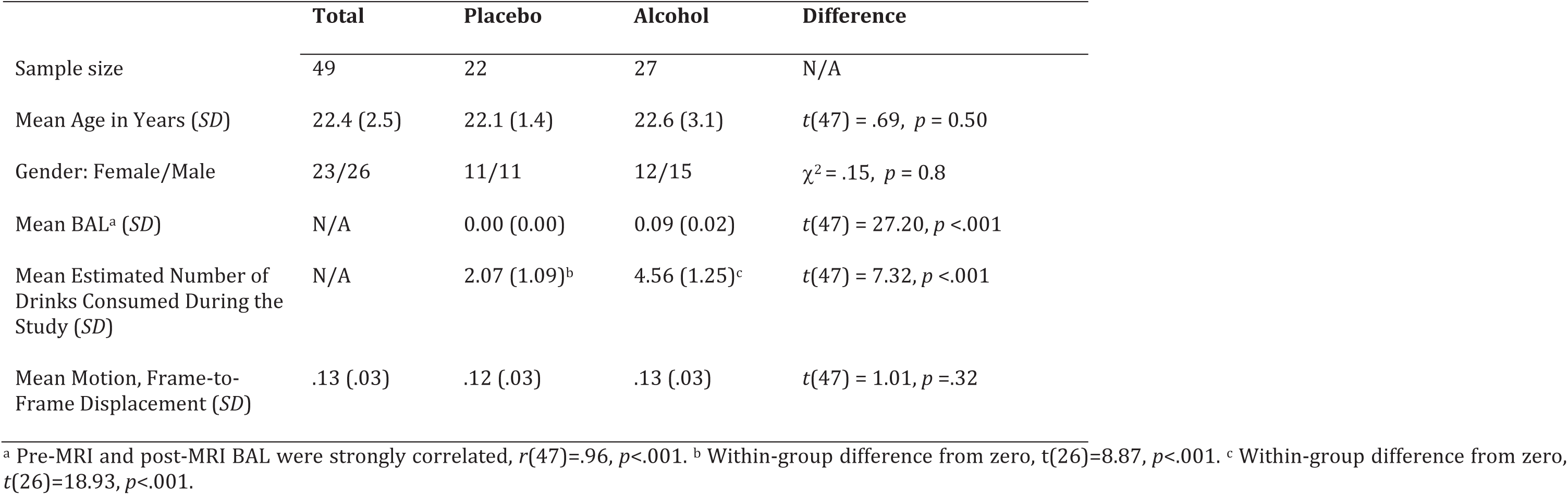
Demographic and descriptive variables.

## METHOD

Methods and materials are summarized below. Detailed descriptions are provided in the Supplement.

### Subjects

Eighty-seven social drinkers (21-3S years old) were recruited from the community as part of a larger study. All reported an absence of substance, neurological, or psychiatric problems. Of these, 61 completed the emotional faces/places paradigm. Twelve subjects were excluded from analyses due to unusable anatomical data (*n*=3), scanner problems (*n*=1), incidental neurological findings (*n*=2), inadequate behavioral performance (>2 *SD*s below the mean; *n*=3), or excessive motion artifact (see below; *n*=3), yielding a final sample of 49 subjects (Table 2). Subjects provided informed written consent. Procedures were approved by the local Institutional Review Board.

### Overview and General Procedures

Subjects abstained from alcohol and other substances for 24 hours and food/drink for 3 hours prior to the session. At the start of the session, subjects were randomly assigned (stratified by sex and race/ethnicity) to receive an alcoholic or placebo beverage, which was consumed just prior to scanning. Blood alcohol level (BAL) was assessed (Alcosensor IV Breathalyzer; Intoximeters Inc., St. Louis, MO) immediately before and after scanning. Subject status was continuously monitored using an MRI-compatible eye-tracker. At the end of the session, subjects estimated the number of standard alcoholic drinks they had consumed.

### Alcohol/Placebo Procedures

Well-established procedures were used for administering alcohol or placebo ^36,37^. Consistent dosing was achieved using a formula that uses height, weight, age, and sex to produce a target BAL of 0.08% or 0.12% with an anticipated variance of ±0.02% ∼30 minutes after the completion of beverage consumption ^38^. This produced a unimodal BAL distribution (range: 0.06% - 0.12%; Table 2). Alcoholic beverages contained a mixture of juice and 100-proof vodka. To control absorption, subjects consumed 3 equal doses over 30 minutes. The placebo group received a similar beverage, with distilled water replacing the vodka. Subjects assigned to the alcohol (or placebo) group observed the experimenter pouring the vodka (or distilled water) from a vodka bottle. The placebo manipulation was reinforced by floating 3 ml of bitters and 3 ml of vodka on the surface of the beverage and delivering a minute amount of aerosolized vodka to the rim of the beverage containers outside the subject’s view. On average, subjects in the placebo group estimated that they consumed ∼2 drinks, validating the manipulation (Table 1).

### Emotional-Faces/Places Paradigm

Building on work by our group ^39^ and others ^19,40^, imaging data were acquired while subjects viewed alternating blocks of emotional faces (8 blocks) or places (9 blocks). Block length (∼16.3 s) was optimized to detect between-condition differences ^41,42^. To minimize habituation ^41–43^, blocks consisted of 16 brief presentations of faces or places (∼1.02 s/image). During face blocks, subjects discriminated threat-related (i.e., fearful; 75% trials) from neutral expressions (25% trials) presented in a pseudorandomized order ^39^. This design choice was aimed at reducing monotony and minimizing potential habituation to the fearful expressions ^43^. During place blocks, subjects discriminated suburban residential residences (i.e., houses; 75%) from urban commercial buildings (i.e., skyscrapers; 25%). Face and place stimuli were adapted from prior work^44,45^.

### MRI Data Acquisition

MRI data were acquired using a Siemens Magnetom TIM Trio 3 Tesla scanner (32-channel head-coil). Sagittal T1-weighted images were acquired using a MPRAGE sequence (TR=1,900 ms; TE=2.32 ms; inversion time=900 ms; flip angle=9°; sagittal slice thickness=0.9 mm; in-plane=0.449×0.449mm; matrix=512×512; field-of-view=230×230). To enhance resolution, a multi-band sequence was used to collect a total of 286 oblique-axial EPI volumes during a single scan of the faces/places task (multiband acceleration=6; TR=1,000 ms; TE=39.4 ms; flip angle=36.4°; slice thickness=2.2 mm, number of slices=60; in-plane resolution=2.1875×2.1875 mm; matrix=96×96). To minimize susceptibility artifacts, images were collected in the oblique axial plane. Co-planar oblique-axial spin echo (SE) images were collected in opposing phase-encoding directions (TR=7,220 ms; TE=73 ms) to enable fieldmap correction.

### MRI Data Preprocessing

MRI data were visually inspected before and after processing for quality assurance.

#### Anatomical Data Processing

Methods are similar to those described in other recent reports by our group ^28,30^. T1 images were brain-extracted (’skull-stripped’) using a multi-tool approach ^30^. Brain-extracted T1 images were normalized to the MNI152 template using the high-precision diffeomorphic approach implemented in *SyN* ^46^. The mean of the normalized T1 images is depicted in **Supplementary Figure S1**. FSL was used to create a fieldmap and undistorted SE image.

#### Functional Data Processing

The first 3 volumes of each EPI scan were removed. Remaining volumes were de-spiked and slice-time corrected using AFNI ^47^. For co-registration of the functional and anatomical images, an average EPI image was created. The average image was simultaneously co-registered with the corresponding T1-weighted image in native space and corrected for geometric distortions using the boundary-based registration method implemented in FSL and the previously created fieldmap, undistorted SE image, and T1 image. Spatial transformations were concatenated and applied to the functional data in a single step. The transformed images were re-sliced (2-mm^3^), smoothed (6-mm), and filtered (0.007812S-Hz high-pass). To assess residual motion artifact, the variance of volume-to-volume displacement of a selected voxel in the center of the brain (*x*=5, *y*=34, *z*=28) was calculated using the motion-corrected EPI data. Subjects (*n*=3) with extreme motion variance (>2*SD*s above the mean) were excluded from analyses.

fMRI data were modeled using SPM12 and in-house MATLAB code. The emotional-faces/places task was modeled using a boxcar function ^48^. Block onsets were modeled using two additional event-related nuisance predictors. Predictors were convolved with a canonical hemodynamic response function. Additional nuisance variates included motion and physiological noise estimates. To attenuate physiological noise, white matter (WM) and cerebrospinal fluid (CSF) time-series were identified by thresholding the tissue prior images distributed with FSL. The EPI time-series was orthogonalized with respect to the first 3 right eigenvectors of the data covariance matrix from the WM and CSF compartments^49^. Reactivity to threat-related faces (i.e., the main effect of Condition: Emotional Faces vs. Places) was assessed using a voxelwise one-sample *t* test controlling for mean-centered age and sex. The impact of alcohol administration was assessed using a voxelwise two-sample *t* test controlling for mean-centered age and sex, equivalent to testing the Group (Alcohol vs. Placebo) x Condition (Emotional Faces vs. Places) interaction.

### Hypothesis Testing Strategy

#### Alcohol-Dampening in the EAc

The first aim of the study was to test the hypothesized dampening effects of acute alcohol administration on EAc reactivity to threat-related faces. Accordingly, the Group x Condition interaction was thresholded at *p*<.05 familywise error (FWE) corrected for the extent of the EAc (**Supplementary Figure S2;** 1,205 voxels; 9,640 mm^3^). The EAc region-of-interest (ROI) encompassed the amygdala, substantia innominata/sublenticular extended amygdala (SI/SLEA), and BST bilaterally ^30,50^. Significant clusters (*p*<.05, whole-brain FWE corrected) outside the EAc are reported on an exploratory basis for voxelwise analyses of the Condition (Emotional Faces vs. Places) and Group x Condition effects.

#### Alcohol-Dampening: BST vs. Ce

The second major aim of our study was to test the differential sensitivity of the BST and the Ce-the two major sub-divisions of the EAc-to the hypothesized threat-dampening effects of alcohol. To do so in an unbiased manner, we extracted and averaged standardized contrast coefficients using anatomically defined, *a priori* ROIs ^21^, as shown in **Supplementary Figure S3**. The BST was defined using the ROI of Theiss and colleagues (2016). The Ce was defined using the ROI of Tillman and colleagues (2018). A mixed-model general linear model was used to compare the impact of Group and Hemisphere on regional reactivity to threat-related faces. Significant interactions were decomposed using the appropriate tests of simple effects. The Group effect is reported using the Welch-Satterthwaite correction for unequal variances (*F_W-S_*).

## RESULTS

### Behavior

On average, subjects were highly accurate at performing the simple discrimination tasks (*M*=86.8%, *SD*=7.9). Nevertheless, performance was ∼8% lower in the alcohol (*M*=83.2%, *SD*=8.2) compared to the placebo group (*M*=91.1%, *SD*=4.9; *F_W-S_*(1,47)=1S.98, *p*<.001), consistent with prior work ^51^. Subjects were ∼4% more accurate when performing the places (*M*=88.8%, *SD*=8.8) compared to the faces discrimination (*M*=84.4%, *SD*=8.4; *F*(48)=22.37, *p*<.001), but the Group x Condition interaction was not reliable (*F*(1,47)=.24, *p*=.63). As noted below and detailed in the Supplement, control analyses indicated that these modest differences in performance were not the primary determinant of alcohol-related differences in neural reactivity.

#### The Dorsal Amygdala is Sensitive to Threat-Related Faces

Within the EAc, threat-related faces were associated with significant activation of the dorsal amygdala, bilaterally (*p*<.05, FWE-corrected; Left: *t*=12.59, volume=1,032 mm^3^; *x*=-20, *y*=-10, *z*=-14; Right: *t*=12.22, volume=1,368 mm^3^; *x*=22, *y*=-8, *z*=-16; Figure 1a and **Supplementary Table 1**), consistent with prior work e.g., ^16^. As shown in **Supplementary Figure S4**, the amygdala cluster overlapped the anatomically defined Ce ROI, with the left and right peaks lying in the dorsocaudal region where the Ce, medial, and basomedial nuclei abut.

**Figure 1.**
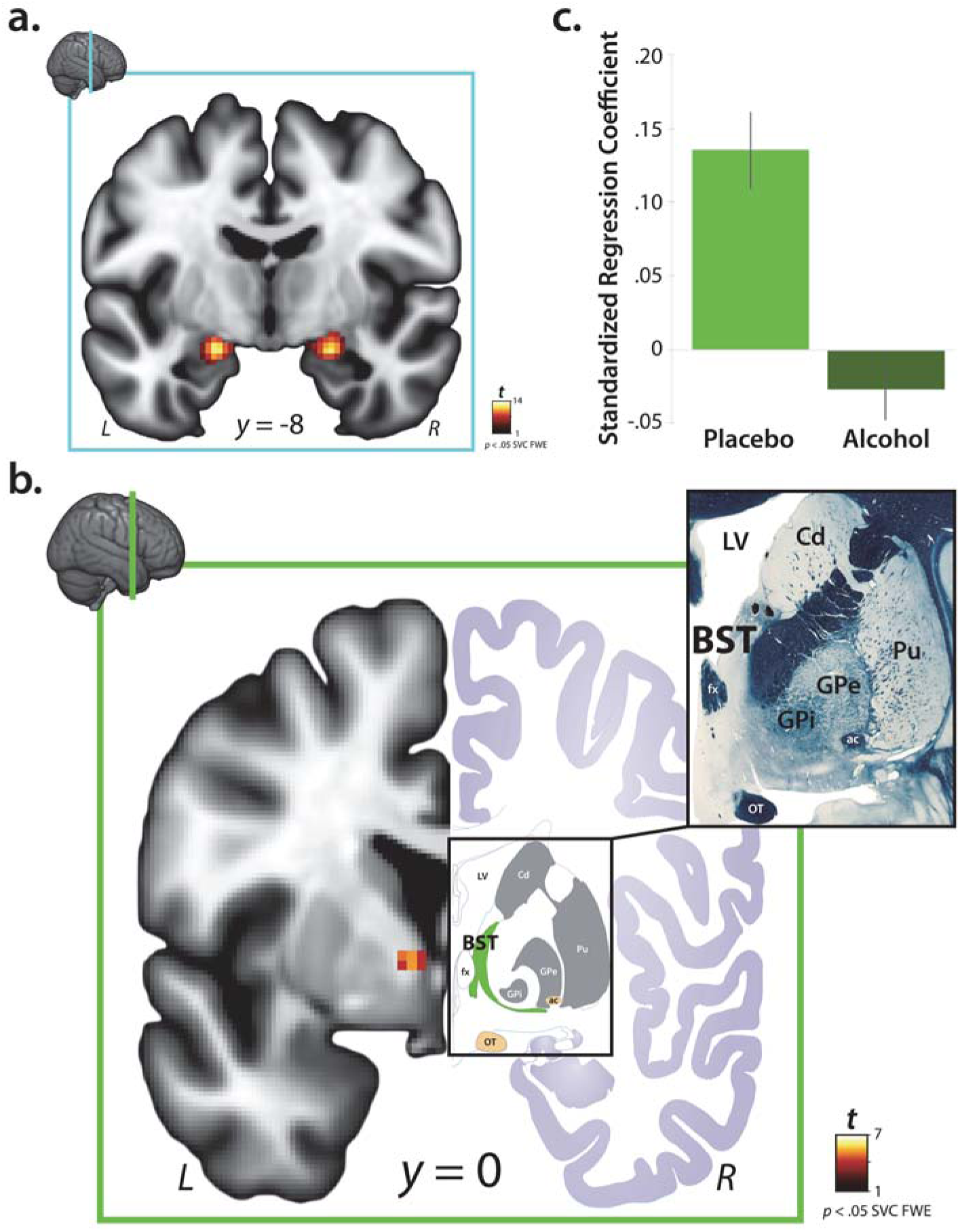
The impact of acute alcohol administration on reactivity to threat-related cues in the central extended amygdala. *A.* Consistent with prior work, voxelwise regression analyses revealed significant activation to threat-related faces in the dorsal amygdala (*p* < .05, FWE corrected for the volume of the anatomically defined EAc region-of-interest; total volume: 1,205 voxels; 9,640 mm^3^). Inset indicates the location of the coronal slice. Significant clusters within the EAc ROI (**Supplementary Figure S2**) are depicted here. For additional results, see **Supplementary Figures S4 and S5** and **Supplementary Tables S1** and **S2**. ***B.*** Voxelwise analyses revealed a significant reduction in reactivity to threat-related faces in the region of the left BST in the alcohol compared to the placebo group (same threshold; equivalent to testing the Group x Condition interaction). The left half of the panel depicts the BST cluster. The right half depicts the BST (*green*) in the corresponding section of the atlas of Mai and colleagues (2015). Note the similar appearance of key landmarks, including the fornix and lateral ventricle (*white*), as well as the optic tract and anterior commissure (*gold*). Upper left inset indicates the location of the coronal slice. Upper right inset depicts the myeloarchitecture (Weigert fiber stain) of this region in the atlas. The left BST was the only significant cluster in EAc-focused or whole-brain analyses. For additional results, see **Supplementary Figure S6** and **Supplementary Table S3**. ***C.*** For illustrative purposes, barplot depicts mean standardized regression coefficients extracted from the peak voxel in the BST cluster for the alcohol (*light green*) and placebo (*dark green*) groups. Hypothesis testing was performed on a voxelwise basis (corrected for multiple comparisons). Error bars indicate the standard error of the mean. Portions of this figure were adapted with permission from the atlas of Mai and colleagues ^54^. Abbreviations-ac, anterior commissure; BST, bed nucleus of the stria terminalis; Cd, caudate; EAc, central division of the extended amygdala; FWE, family-wise error; fx, fornix; GPe, external globus pallidus; GPi, internal globus pallidus; L, left hemisphere; LV, lateral ventricle; OT, optic tract; Pu, putamen; R, right hemisphere; SVC, small volume correction.

On an exploratory basis, we also computed a series of whole-brain analyses. Results indicated that the dorsal amygdala and fusiform cortex (’fusiform face area’) were significantly more sensitive to threat-related faces, whereas the parahippocampal cortex (’parahippocampal place area’) was significantly more sensitive to places, as expected ^52,53^(*p*<.05, FWE-corrected; **Supplementary Figure S5** and **Supplementary Table 2**).

#### Alcohol Dampens BST Reactivity to Threat-Related Faces

Within the EAc, acute alcohol administration was associated with a significant reduction in left BST reactivity to threat-related faces (Group x Condition: *p*<.05, FWE-corrected; *t*=5.46, volume=104 mm^3^; *x*=-8, *y*=-2, *z*=0; Figures 1b-1c and **Supplementary Table 3**). As shown in **Supplementary Figure S6**, the left BST cluster overlapped the anatomically defined BST ROI. Exploratory whole-brain analyses revealed no additional clusters. As detailed in the Supplement, control analyses indicated that the dampening effects of alcohol on BST reactivity to threat-related faces were not a consequence of group differences in performance.

#### Alcohol Exerts Similar Effects in the Ce and the BST

To assess regional differences in EAc activation in an unbiased manner ^21^, standardized contrast coefficients (i.e., faces vs. places) were extracted from the left and right Ce and BST-the two major subdivisions of the EAc-using anatomically defined, *a priori* ROIs, as shown in the upper portion of Figure 2 (Ce: *cyan*; BST: *green*). A mixed-model GLM was then used to compare the impact of Group and Hemisphere on regional reactivity to threat-related faces. Analyses revealed greater threat-related activation in the Ce relative to the BST (Region: *F*(1,47)=32.99, *p*<.001) and a near-significant alcohol-dampening effect across regions (Group: *F_W-S_*(1,47)=3.93, *p*=.053]. Other omnibus effects were not significant (*p*s>.15). Analyses performed using a performance-matched sub-sample revealed similar results (**Supplement**). Collectively, these observations indicate that alcohol acutely dampens reactivity to threat-related faces and it does so similarly in the Ce and the BST.

**Figure 2.**
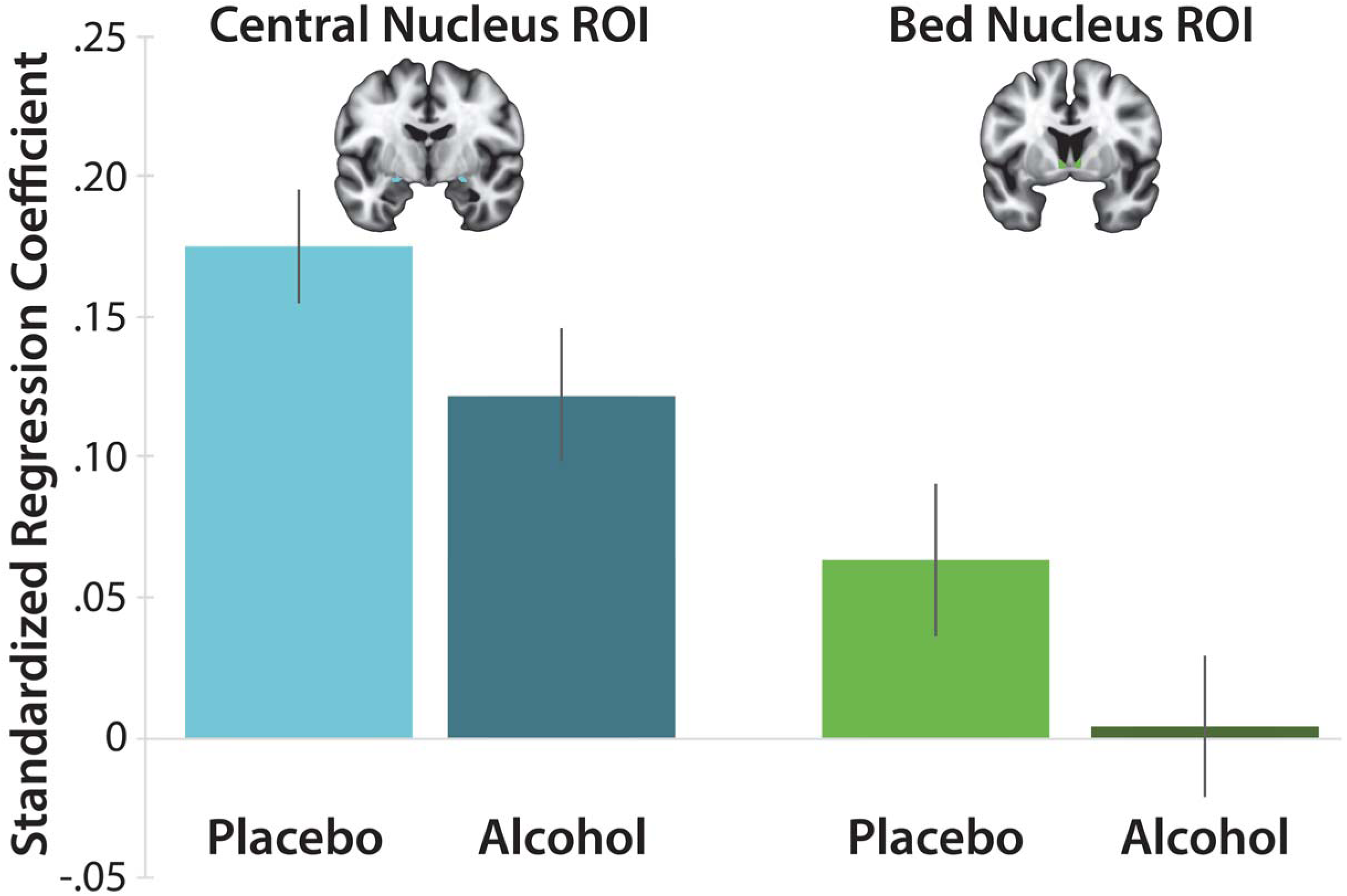
The impact of acute alcohol administration on the two major divisions of the EAc. Barplot depicts mean regression coefficients associated with the emotional-faces/places task for the anatomically defined Ce and BST ROIs for each group. The Ce was significantly more reactive to threat-related faces, relative to the BST (*p*<.001). On average, subjects randomly assigned to the alcohol group showed significantly less reactivity to threat-related faces, relative to those in the placebo group (*p* =.053; equivalent to testing the Group x Condition interaction). The Group x Region interaction was not significant (*p* =.88), suggesting that the Ce and BST are similarly sensitive to the threat-dampening impact of acute alcohol administration. Error bars indicate the standard error of the mean. Abbreviations-EAc, central extended amygdala; ROI, region of interest.

## DISCUSSION

Recent epidemiological work indicates that “the United States is facing a crisis with alcohol use, one that is currently costly and about to get worse” ^55^, yet the neural circuitry most relevant to human alcohol consumption has remained incompletely understood. The etiology of alcohol abuse is complex and involves multiple neurocognitive and motivational systems ^5^, but observations gleaned from clinical research in humans and mechanistic work in rodents highlights the potential importance of negative reinforcement effects mediated by the EAc ^5,8^. Leveraging a placebo-controlled randomized-groups design, the present results demonstrate for the first time that that acute alcohol administration significantly dampens reactivity to threat-related faces in the BST (Figure 1). Analyses performed using unbiased, anatomically defined ROIs revealed a similar pattern of alcohol dampening across the Ce and BST (Figure 2). Control analyses indicated that these results were not an artifact of group differences in performance (**Supplement**). Collectively, these findings indicate that acute alcohol intoxication dampens reactivity to threat-related faces in humans and it does so similarly across the EAc.

The present results reinforce the translational relevance of addiction models derived from preclinical research in mice and rats ^5,9^. Work in rodent models directly implicates the EAc in the threat-dampening consequences of alcohol ^7,56–59^. Immediate early gene studies show that alcohol robustly engages both the Ce and the BST ^60^. While the molecular consequences of alcohol are complex, acute alcohol inhibits excitatory (i.e., glutamatergic) neurotransmission in the Ce and the BST ^61–63^ and increases the release of the inhibitory neurotransmitter gamma-aminobutyric acid (GABA) in the Ce via interactions with the corticotropin-releasing factor (CRF) type-1 receptor ^62,64–69^. Enhanced GABAergic tone within the Ce is, in turn, thought to inhibit cells in the BST and other downstream effector regions ^58,65^. Other work indicates that CRF projections from the Ce to the BST play a critical role in excessive drinking in alcohol-dependent rats ^70^, consistent with evidence implicating the EAc in withdrawal-related negative affect and stress-induced substance use ^31^. While our observations align with this body of research, as with any human neuroimaging study, our conclusions are tempered by questions about the origins and significance of the blood oxygen level-dependent fMRI signal ^71^.

The present results are broadly consistent with clinical research underscoring the importance of negative reinforcement mechanisms in recreational drinking as well as alcohol abuse. Many drinkers expect alcohol to reduce tension or stress ^72^ and those seeking stress reduction are at greater risk for developing an AUD ^73,74^. Individuals with a more anxious temperament and patients with anxiety orders (e.g., social phobia) are more likely to misuse alcohol ^75,76^, and homologous effects have been found in rodents ^6,77^. In the laboratory, moderate doses of alcohol selectively dampen defensive responses (e.g., startle) elicited by uncertain physical danger ^8^. Our results suggest that some of these effects may reflect a downstream consequence of dampened EAc reactivity to potential threat.

Although static images of fearful faces do not elicit robust signs of fear or anxiety ^78^, they are relevant to daily experience, can increase anxiety symptoms, and are perceived as more threatening and arousing than neutral or happy faces ^79–81^. Fearful faces have also been shown to promote vigilance, increasing visual sensitivity, boosting the resolution of visual processing, and enhancing the efficiency of attentional search ^16^. Vigilance is thought to be mediated by circuits centered on the EAc and, once elicited, increases the likelihood experiencing more extreme or pervasive states of anxiety ^16,82^. A key challenge for future research will be to examine the impact of alcohol on EAc reactivity to more intense anxiety-provoking stimuli, such as uncertain threat-of-shock. This approach would dovetail with work in rodent models, enhancing the likelihood of successful bi-directional translation ^34^. Prospective longitudinal imaging research would be useful for understanding the relevance of EAc circuitry to the development of alcohol use disorder and other addictions. Combined with more naturalistic measures of stress-induced drinking in the laboratory or field (e.g., ecological momentary assessment), this approach might provide a means of stratifying at-risk populations or patients into the subset for whom negative reinforcement circuits are most relevant to intervention.

The present results are also relevant to on-going debates about the functional organization of the EAc ^13^. Among researchers focused on humans, it is widely believed that the Ce and BST are functionally dissociable. Inspired by an earlier generation of lesion and inactivation studies in rodents ^83^, this ‘double-dissociation’ or ‘strict-segregation’ hypothesis suggests that the Ce (or the amygdala more generally) rapidly assembles phasic responses to clear-and-immediate threats (e.g., a cue associated with the imminent delivery of shock), whereas the BST comes on-line more slowly and is responsible for orchestrating sustained responses to dangers that are diffuse, uncertain, or remote. While this hypothesis remains popular, and has even been incorporated into the National Institute of Mental Health’s Research Domain Criteria (RDoC) initiative, a range of evidence gleaned from studies of rodents, monkeys, and humans makes it clear that while the Ce and the BST are certainly not interchangeable, they are more alike than different ^12,84^. Leveraging an unbiased ROI approach, our results extend this work to show that the Ce and BST are similarly sensitive to the threat-dampening effects of alcohol. Whether this generalizes to more intense threat-related cues (e.g., film clips) or other anxiolytics (e.g., benzodiazepines) remains an important avenue for future studies.

Existing treatments for alcohol use and other additions are far from curative ^4,85^, highlighting the need to develop a deeper understanding of the underlying motivational processes and neural systems. The present results demonstrate that acute alcohol intoxication dampens reactivity to threat-related cues across the human EAc. The use of a relatively large sample (Table 1), placebo-controlled between-groups design, ecologically relevant dosing, fMRI-optimized task, best practices for the acquisition and processing of functional neuroimaging data, and unbiased ROI analytic approach enhances confidence in the clinical and translational significance of these results. More broadly, these findings set the stage for mechanistic work aimed at developing more effective treatments for alcohol abuse and other debilitating addictions.

## CONTRIBUTIONS

C.M.K, A.J.S., and J.F.S. designed the imaging study based on procedures originally developed and refined by D.E.B. and J.J.C. for psychophysiological research. C.M.K. and J.F.S. collected data. J.H., C.M.K., and J.F.S. processed data. J.H., C.M.K., A.J.S., and J.F.S. analyzed data. J.H., A.J.S., A.S.F., and J.F.S. interpreted data. J.H., C.M.K., J.F.S., and A.J.S. wrote the paper. A.J.S. created figures. J.H., J.F.S., and A.J.S. created tables. A.S.F., D.E.B., and J.J.C. provided theoretical guidance. A.J.S. funded and supervised all aspects of the study. All authors contributed to reviewing and revising the paper and approved the final version.

## ACKNOWLEDGEMENTS

Authors acknowledge critical feedback from E. Bernat, L. Pessoa, and M. Roesch and assistance from K. DeYoung, L. Friedman, M. Gamer, B. Nacewicz, M. Stockbridge, S. Padmala, P. Spechler, R. Tillman, and personnel from the Affective and Translational Neuroscience Laboratory and Maryland Neuroimaging Center. This work was supported by the University of California, Davis; University of Maryland, College Park; University of Wisconsin-Madison; and National Institutes of Health (DA040717 and MH107444). Development of the NimStim Face Database was overseen by N. Tottenham and supported by the MacArthur Foundation. Authors declare no conflicts of interest.

## DATA AVAILABILITY/SHARING

All of the key statistical maps and regions-of-interest have been or will be uploaded to NeuroVault.org.

