## Supplementary Materials for "Acute alcohol administration dampens threat-related activation in the central extended amygdala"

***Contents***

| | |
|---|---|
| Supplementary Comment on the Term 'Threat-Related' | Pages 3 |
| Supplementary Method | Pages 3-13 |
| Supplementary Figure S1. Mean normalized T1 image. | Page 14 |
| Supplementary Figure S2. EAc ROI. | Page 15 |
| Supplementary Figure S3. BST and Ce ROIs. | Page 16 |
| Supplementary Figure S4. The amygdala cluster identified in voxelwise analyses overlaps the anatomically defined Ce ROI. | Page 17 |
| Supplementary Table 1. Descriptive statistics for clusters identified by the faces vs. places contrast… | |
| using p<.05, small-volume corrected | Pages 18-22 |
| Supplementary Figure S5. Regions identified by a whole-brain, voxelwise regression analysis | Page 23 |
| Supplementary Table 2. Descriptive statistics for clusters identified by the faces vs. places contrast… | |
| using p<.05, whole-brain corrected | Pages 24-26 |
| Supplementary Figure S6. The BST cluster identified in voxelwise analyses overlaps the anatomically defined BST ROI | Page 27 |
| Supplementary Table 3. Descriptive statistics for clusters identified by the Group × Condition contrast… | |
| using p<.05, small-volume corrected | Page 28 |
| Analyses Controlling for Group Differences in Performance | Pages 29-30 |
| Supplementary Table 4: Mean Percentage Correct for the Complete Sample | Page 29 |
| Supplementary Table 5: Mean Percentage Correct for the Performance-Matched Sub-Sample | Page 29 |
| Supplementary References | Pages 30-39 |

**Supplementary Comment on the Term 'Threat-Related'**

As we have previously noted [1,2], the term 'threat-related' encompasses a broad range of stimuli, including clear and immediate dangers (e.g., cues paired with shock), novel situations or individuals, uncertain or diffuse dangers (e.g., darkness), aversive stimuli (e.g., unpleasant images), and angry and fearful facial expressions. Fearful faces, in particular, signal the presence, but not the source of potential threat, and promote heightened vigilance in the absence of defensive mobilization [3]. That is, static images of fearful faces do not amplify the startle reflex [4,5] or autonomic measures [6]. But they can increase subjective feelings of anxiety [7] and are perceived as more threatening and arousing than neutral or happy faces [5,8]. They also increase vigilance for potentially threat-relevant information. Fearful faces have been shown to increase contrast sensitivity [9] and orientation sensitivity [10], to boost the spatial and temporal resolution of visual processing [10], and to enhance the efficiency of visual search [11].

### SUPPLEMENTARY METHOD

#### *Subjects*

A total of 61 individuals between the ages of 21 and 35 years were recruited from the community as part of a larger study. All had experience with the highest study dose of alcohol used in the present study (~4-5 standard drinks) within the past 12 months, normal or corrected-to-normal color vision, and reported the absence of lifetime alcohol or substance-related problems, lifetime neurological symptoms, current psychiatric diagnosis or treatment, pervasive developmental disorder or very premature birth, or a medical condition that would contraindicate either acute alcohol consumption or MRI. Twelve subjects were excluded from analyses due to unusable T1-weighted datasets ($n$=3), technical problems with the scanner ($n$=1), incidental neurological findings ($n$=2), inadequate behavioral performance (>2 *SD*s below the mean; $n$=3), or excessive motion artifact ($n$=3; see below), yielding a final sample of 49 subjects (46.9% female; **Table 2** in the main report). Subjects provided informed written consent and all procedures were approved by the University of Maryland Institutional Review Board.

#### *Overview and General Procedures*

Subjects were instructed to abstain from alcohol and other drugs for at least 24 hours and from all food and drink for at least 3 hours prior to the experimental session. At the start of the session, subjects were randomly assigned (stratified by sex and race/ethnicity) to receive either an alcoholic or a placebo beverage, inclusionary/exclusionary criteria (e.g., MRI contraindications) were re-assessed, and initial sobriety was confirmed using a standard breath assay (Alcosensor IV Breathalyzer; Intoximeters Inc., St. Louis, MO). Both groups were informed that they could receive a moderately impairing dose of alcohol. After obtaining consent, an overview of the experimental procedures was provided and the emotional-faces/places task was described (see below). Just prior to scanning, subjects consumed an alcoholic or placebo beverage using well-established procedures (see below). Noise-dampening, MRI-compatible ear buds were provided for communication and hearing protection. Before entering the scanner room, a handheld metal detector was used to verify the absence of potentially hazardous ferromagnetic objects. The subject was positioned prone in the scanner and foam inserts were used to minimize potential movement. Visual stimuli were digitally projected onto a screen mounted at the head-end of the scanner bore and viewed using a mirror mounted on the head-coil. Subject status was continuously monitored from the control room using an MRI-compatible eye-tracker (data not recorded; Eyelink 1000; SR Research, Ottawa, Ontario, Canada). Head motion was monitored in real time using AFNI [12]. Blood alcohol level (BAL) was assessed via breath assay in the control room immediately before and after the imaging component of the session. Following the post-MRI breath assay, subjects were asked to estimate the number of standard alcoholic drinks consumed during the session. Subjects were then debriefed and compensated. Those assigned to the Alcohol group were required to remain at the imaging center until their estimated BAL was <0.03%.

### Alcohol/Placebo Procedures

Procedures for administering and quantifying acute alcohol administration were adapted from techniques established by Curtin and colleagues for psychophysiological research [13,14] and are similar to those used in prior imaging research [15]. Dosing was titrated using a formula that uses height, weight, age, and sex to produce a target BAL of 0.08% or 0.12% with an anticipated variance of ±0.02% ~30 minutes after the completion of beverage consumption [16]. This produced a unimodal distribution of mean BAL (range: 0.061% - 0.119%; see **Table 2** in the main report). Alcoholic beverages contained a mixture of cranberry juice, Tang™, and 100-proof vodka (Smirnoff Blue™). To control absorption rates, subjects consumed 3 equal doses, evenly spaced over 30 minutes. Subjects in the placebo control group received a similar beverage, with an equivalent volume of distilled water replacing the vodka. Subjects assigned to the alcohol (or placebo) group observed the experimenter pouring the vodka (or distilled water) from a vodka bottle. The placebo manipulation was reinforced by floating 3 ml of bitters and 3 ml of vodka on the surface of the beverage and delivering a minute amount of aerosolized vodka to the rim of the beverage containers (i.e., in an adjacent room, outside of the subject's view). Immediately following consumption of the third beverage, BAL was assessed and subjects were scanned. BAL was re-assessed immediately following the final scan (inter-assessment period: $M$ = 70 min, $SD$ = 6.0 min) see [17,18].

On average, subjects assigned to the placebo group estimated that they consumed ~2 drinks, confirming the efficacy of the placebo manipulation (**Table 1**). All subjects tolerated the beverages without complications or adverse effects.

**Emotional-Faces/Places Paradigm**

To assess the impact of acute alcohol administration on threat-related reactivity in the EAc, imaging data were acquired while subjects performed a simple, fMRI-optimized, continuous-performance task. Building on work by our group [2,19] and others [17,20-22] demonstrating the utility of emotional face paradigms for probing amygdala reactivity, subjects viewed alternating blocks of either emotional faces (8 blocks) or places (9 blocks). The use of a block design mitigates potential concerns about alcohol-induced changes in the shape of the hemodynamic response function (HRF). Block length (~16.3 s) was chosen to maximize our power to detect a difference in the blood oxygen level-dependent (BOLD) signal elicited by the two conditions [23,24]. To maximize signal strength and homogeneity and minimize potential neural habituation [23-25], each block consisted of 16 brief presentations of faces or places (~1.02 s/image) (for related approaches, see Williams et al., 2015 and O'Craven & Kanwisher, 2000). During face blocks, subjects used an MRI-compatible, fiber-optic response pad (MRA, Washington, PA) to discriminate (i.e., two-alternative, forced choice) between threat-related (i.e., fearful; 75% trials) and emotionally neutral expressions (25% trials) presented in a pseudorandomized order [2,3,26,27]. This design choice was aimed at reducing monotony and minimizing potential habituation to the fearful expressions [25]. During place blocks, subjects discriminated between suburban residential buildings (i.e., houses; 75%) and urban commercial buildings (i.e., skyscrapers; 25%). Face stimuli were taken from prior work by Gamer and colleagues [28,29] and included standardized images of unfamiliar male and female adults displaying unambiguous fearful or neutral expressions. To maximize the number of models and mitigate potential habituation, images were derived from several well-established databases: Ekman and Friesen's

Pictures of Facial Affect [30], the FACES database [31], the Karolinska Directed Emotional Faces database (http://www.emotionlab.se/resources/kdef), and the NimStim Face Stimulus Set (https://www.macbrain.org/resources.htm). Colored images were converted to grayscale, brightness normalized, and masked to occlude non-facial features (e.g., ears, hair). Grayscale building stimuli were taken from prior work by Choi and colleagues [32,33].

**MRI Data Acquisition**

MRI data were acquired using a Siemens Magnetom TIM Trio 3 Tesla scanner and 32-channel head-coil. Sagittal T1-weighted anatomical images were acquired using a magnetization-prepared, rapid-acquisition, gradient-echo (MPRAGE) sequence (TR=1,900 ms; TE=2.32 ms; inversion time=900 ms; flip angle=9°; sagittal slice thickness=0.9 mm; in-plane=0.449 × 0.449mm; matrix=512 × 512; field-of-view=230 × 230). To enhance spatial and temporal resolution, a multi-band sequence was used to collect a total of 286 oblique-axial echo planar imaging (EPI) volumes during a single scan of the faces/places task (multiband acceleration=6; TR=1,000 ms; TE=39.4 ms; flip angle=36.4°; slice thickness=2.2 mm, number of slices=60; in-plane resolution=2.1875 × 2.1875 mm; matrix=96 × 96). Images were collected in the oblique axial plane (approximately -20° relative to the AC-PC plane) to minimize susceptibility artifacts. To enable fieldmap correction, two oblique-axial spin echo (SE) images were collected in each of two opposing phase-encoding directions (rostral-to-caudal and caudal-to-rostral) at the same location and resolution as the functional volumes (i.e., co-planar; TR=7,220 ms; TE=73 ms).

**MRI Data Preprocessing**

Given our focus on the EAc, methods were optimized to minimize spatial normalization error and other kinds of noise.

***Anatomical Data Processing.*** Methods are similar to those described in recent reports by our group [19,34] and others [35,36] and are only summarized here. In brief, prior work indicates that spatial normalization is enhanced by using a brain-extracted (i.e., 'skull-stripped') template and brain-extracted T1-weighted 'anatomical' images [37-39]. To ensure consistently high-quality extractions, we implemented a multi-tool strategy. T1-weighted images were written into standard orientation, re-scaled (0-1,000) using ANTs' *ImageMath* tool and inhomogeneity-corrected using N4 [40]. Four extraction masks were generated for each T1-weighted image. Three masks were generated using *BSE* [41], *ROBEX* [42], and SPM unified segmentation [43] with an updated tissue probability map [44], respectively. The fourth mask was generated by spatially normalizing the full T1 volume (unextracted) to the MNI152 template (ICBM 152 non-linear 6th-generation asymmetric average brain stereotaxic registration model) [45] using the high-precision diffeomorphic approach implemented in *SyN* (mutual information cost function) [42,46-48]. The inverse of this transformation was then applied to a manually refined version of the 1-mm MNI152 brain mask distributed with FSL. Next, a best-estimate extraction mask was determined by consensus, requiring agreement across three or more extraction masks [49-52]. Using this consensus mask, each T1-weighted image was extracted and spatially normalized to a manually refined, brain-extracted version of the 1-mm MNI152 template using *SyN*. Brain-extracted T1 images were segmented using *FAST* [53] for subsequent use in EPI-to-T1 co-registration (see below). Each dataset was visually inspected before and after processing for quality assurance. Brain extraction failed for one subject. In this case, a mask was manually generated using a combination of SPM and *SyN* (see above) as well as *BET2* [54] and AFNI's *3dSkullStrip* (Cox, 1996). The mean of the 49 spatially normalized T1-weighted images is depicted in **Supplementary Figure S1**.

***Fieldmap Processing*.** Co-planar SE images with opposing phase-encoding directions were written to standard orientation and *topup* (https://fsl.fmrib.ox.ac.uk/fsl/fslwiki/topup) was used to create a fieldmap and undistorted SE image [55,56]. To mitigate extreme values, fieldmaps were converted to radians; de-spiked and median filtered using default parameters; thresholded (min=-375, max=475); and spatially smoothed (smooth2=5, smooth3=2) using a combination of *fugue* and *fslmaths*. The average of the undistorted SE magnitude images was inhomogeneity corrected using *N4* and brain extracted using *3dSkullStrip*.

***Functional Data Processing***. The first 3 volumes of each EPI scan were removed to allow for signal saturation. Remaining volumes were written to standard orientation using FSL and then de-spiked (*3ddespike*) and slice-time corrected (*3dtshift*) using default settings in AFNI [12]. Recent methodological work indicates that de-spiking is more effective than 'scrubbing' [57-60] for attenuating motion-related artifacts. Slice-time correction was performed using the multiband acquisition times and shifting each slice to the midpoint of the volume acquisition. For co-registration of the functional and anatomical images, an average EPI image was created using AFNI's *3dvolreg* two-pass motion correction. The average image was simultaneously co-registered with the corresponding T1-weighted image in native space and corrected for geometric distortions using FSL's *EPI_reg.sh* with the boundary-based cost function and the previously created fieldmap, undistorted average SE image, inhomogeneity-corrected T1-weighted image, and white matter compartment. Unpublished observations by our group indicate that, even after spatially smoothing and thresholding the fieldmap, the resulting distortion-corrected EPI images often contain areas that appear 'over-corrected,' particularly in midline regions of the medial temporal lobe and orbitofrontal cortex. In order to explicitly control the smoothness of the distortion correction as well as to create an ITK-compatible version of the distortion correction warp field, we used ANTs to spatially transform the average EPI image (i.e., prior to processing with *EPI_reg.sh*) directly to the co-registered, distortion-corrected image output from

*EPI_reg*. The time-series images were then motion-corrected by rigidly aligning each volume to the average image using *AntsMotionCorr* and the estimated affine transformation matrix was saved for each image. At this point, the complete set of spatial transformations for each volume is known and was applied in a single step. Thus, for each slice-time corrected image, the following spatial transformations were concatenated and applied: motion-correction rigid-body, co-registration-unwarping affine, co-registration-unwarping nonlinear, brain-extracted T1 affine, brain-extracted T1 diffeomorphic. The transformed images were re-sliced to 2-mm$^3$ (5$^{th}$-order splines), spatially smoothed (6-mm FWHM) within the brain mask using AFNI's *3dBlurInMask*, and temporally filtered (0.0078125-Hz high-pass) using SPM12 (http://www.fil.ion.ucl.ac.uk/spm). To assess residual motion artifact, the variance of volume-to-volume displacement of a selected voxel in the center of the brain ($x$=5, $y$=34, $z$=28) was calculated using the motion-corrected EPI data. Subjects ($n$=3) with extreme motion variance (>2*SD*s above the mean) were excluded from analyses.

**fMRI Modeling**

fMRI data were modeled using SPM12 and in-house MATLAB code. At the first level (single-subject), the emotional-faces/places task was modeled using a simple boxcar function with place blocks serving as the implicit baseline [61]. Nearly identical results, with opposite sign, were observed when face blocks were treated as the implicit baseline (not reported). Nuisance variance attributable to block onsets (e.g., set-switching processes) was modeled using two additional event-related predictors. All three predictors were convolved with a canonical HRF. Additional nuisance variates included 19 estimates of motion (rostral-caudal, dorsal-ventral, left-right, pitch, roll, and yaw lagged by 0, 1, and 2 volumes; and the final value of the cost function minimized during rigid-body motion correction [negative mutual information with the mean EPI image]) was also included as a regressor of no interest. ) and 2 estimates of physiological noise. To attenuate physiological noise, white matter (WM) and cerebrospinal fluid (CSF) time-series were identified by thresholding the tissue prior images distributed with FSL, as in prior work by our group [62,63] and others [64]. The EPI time-series was orthogonalized with respect to the first 3 right eigenvectors of the data covariance matrix from the WM and CSF compartments [65]. Prior research in relatively large samples (*n*=88: 0.06% BAL vs. Placebo) has failed to uncover alcohol-induced changes in EAc blood flow, mitigating concerns about gross hemodynamic differences [66].

Reactivity to threat-related faces (i.e., the main effect of Condition: Emotional Faces vs. Places) was assessed using a voxelwise one-sample *t* test controlling for nuisance variance in mean-centered age and sex. The impact of alcohol administration was assessed using a voxelwise two-sample *t* test controlling for mean-centered age and sex, equivalent to testing the Group (Alcohol vs. Placebo) × Condition (Emotional Faces vs. Places) interaction. The effects of alcohol were tested on both the [Emotional Faces > Places] and [Places > Emotional Faces] contrasts.

### Hypothesis Testing Strategy

***Alcohol-Dampening in the EAc.*** The first major aim of the present study was to test the hypothesized dampening effects of acute alcohol administration on EAc reactivity to threat-related faces. Accordingly, the Group × Condition interaction (i.e., voxelwise two-sample *t* test) was thresholded at $p<.05$ familywise error (FWE) corrected for the extent of the EAc (**Supplementary Figure S2**), as in prior work by our group [1]. The EAc region-of-interest (ROI) encompassed the amygdala, substantia innominata/sublenticular extended amygdala (SI/SLEA), and BST bilaterally [63,67]. Consistent with recent recommendations [68,69], the EAc ROI was created using a combination of the Mai and Harvard-Oxford atlases [70-74] and included the probabilistic BST ROI developed by Theiss and colleagues (*p*>0%) [75] and the Harvard-Oxford probabilistic amygdala. Using this as a starting point, voxels in the region of the SI/SLEA was manually added in the coronal plane of the 1-mm MNI152 template, working from rostral to caudal, and confirmed in the other planes. At intermediate levels of the amygdala's rostral-caudal axis, where the BST was no longer visible, the SI/SLEA was limited to voxels dorsal to the amygdala and ventral to the putamen and pallidum. SI/SLEA voxels were included until the head of the hippocampus was clearly visible. Voxels in neighboring regions of the accumbens, caudate, putamen, pallidum, thalamus, and ventricles (Harvard-Oxford atlas, *p*>50%) were excluded using a Boolean 'NOT.' The resulting bilateral ROI was decimated to 2-mm$^3$ (total: 1,205 voxels; 9,640 mm$^3$). Significant clusters (*p*<.05, whole-brain FWE corrected) outside the EAc are reported on an exploratory basis for voxelwise analyses of Condition (i.e., one-sample *t* test of Emotional Faces vs. Places) and the Group × Condition interaction (i.e., two-sample *t* test). Clusters were labeled using a combination of the Mai and Harvard-Oxford atlases and related work [67,76]. Figures were created using MRIcron (http://people.cas.sc.edu/rorden/mricron) and MRIcroGL (http://www.mccauslandcenter.sc.edu/mricrogl).

***Alcohol-Dampening: BST vs. Ce.*** The second major aim of our study was to test the differential sensitivity of the BST and the Ce—the two major sub-divisions of the EAc—to the hypothesized threat-dampening effects of alcohol. To do so in an unbiased manner, we extracted and averaged standardized contrast coefficients (i.e., beta weights) using anatomically defined, *a priori* ROIs [77], as shown in **Supplementary Figure S3**. The BST was again defined using the probabilistic ROI of Theiss and colleagues and the Ce was defined using the ROI described by Tillman and colleagues (2018). A mixed-model general linear model (GLM) was then used to compare the impact of Group and Hemisphere on regional reactivity to threat-related faces. Significant interactions were decomposed using the appropriate tests of simple effects. The main effect of Group is reported using the Welch-Satterthwaite correction for unequal variances ($F_{W\text{-}S}$). ROI and behavioral analyses were computed using SPSS (version 24.0.0.0; IBM Inc., Armonk, NY). Ancillary analyses indicated that the data adequately satisfied GLM assumptions [78,79]. We use the terms more and less 'activated' to refer to relative differences in the blood oxygen level-dependent (BOLD) signal for particular task contrasts, as indexed by standardized regression coefficients. Naturally, relative differences in 'activation' reflect more complex relations between hemodynamic and different kinds of cellular activity [80].

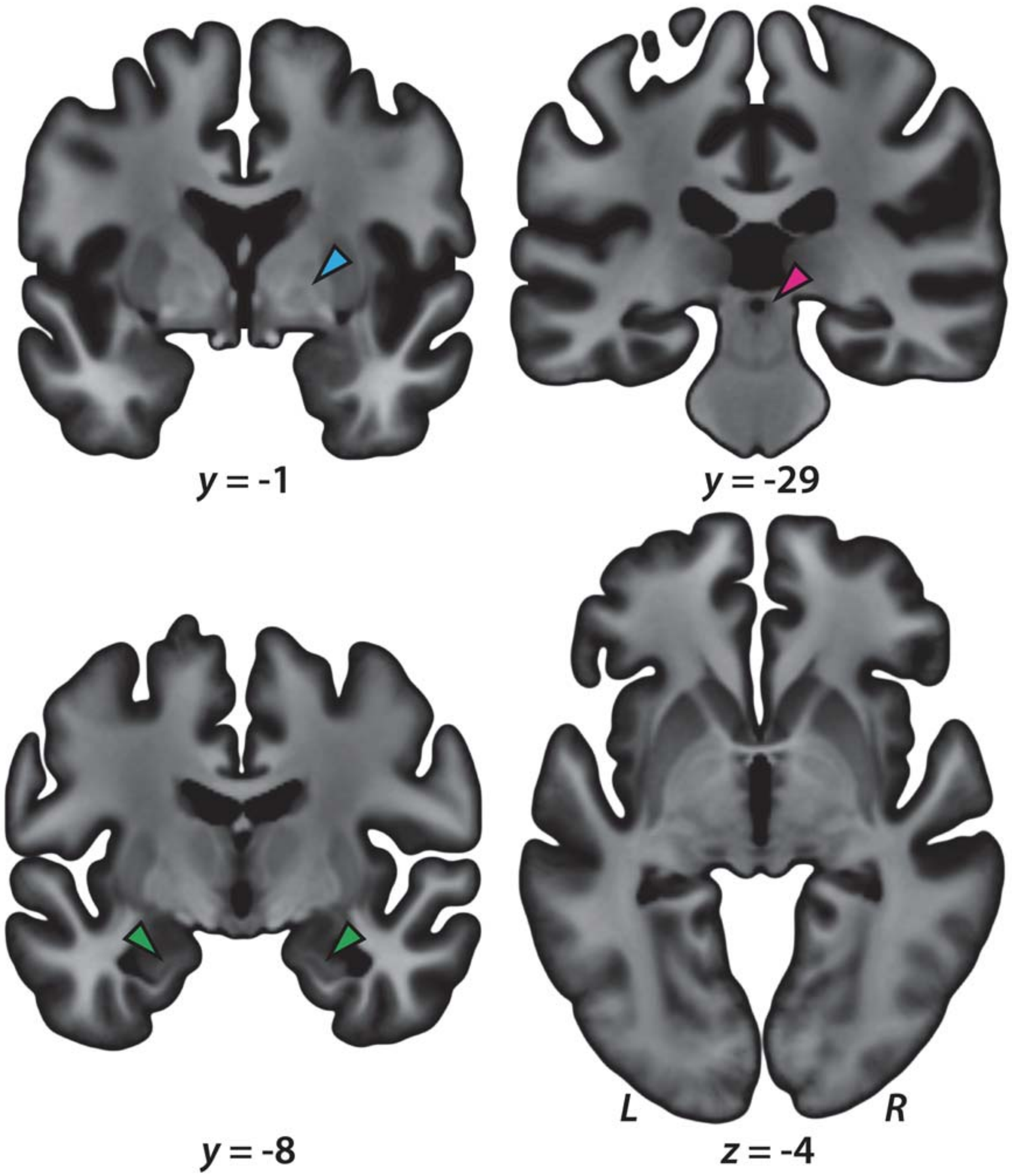

**Supplementary Figure S1.** ***Mean normalized T1 image***. Figure depicts representative slices from the average of the 49 diffeomorphically normalized T1-weighted images. Note the preservation of fine detail in the medial medullary lamina of the globus pallidus (*cyan* arrowhead), periaqueductal gray (*magenta* arrowhead), and alveus (*green* arrowheads).

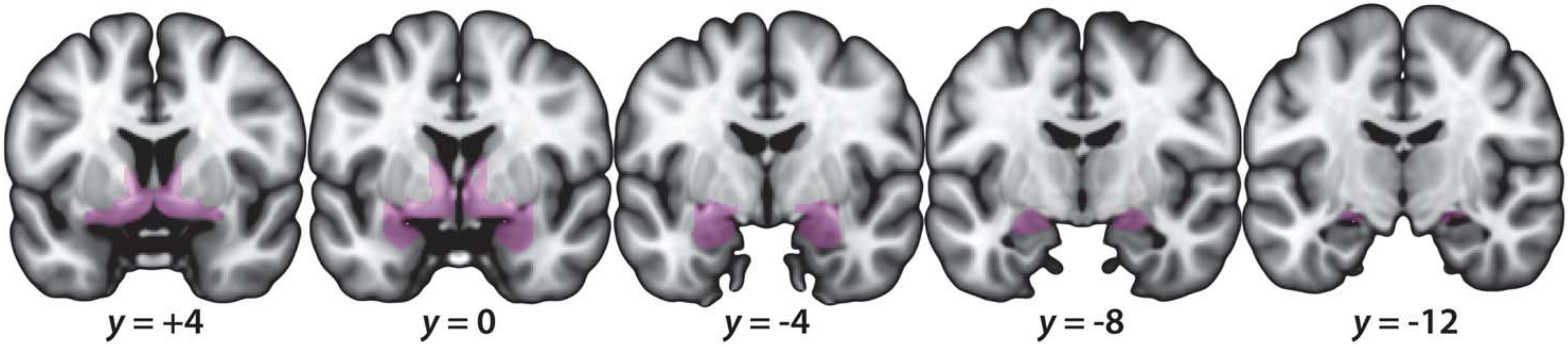

**Supplementary Figure S2. EAc ROI.** The EAc ROI (*purple*) encompassed the amygdala, substantia innominata/sublenticular extended amygdala (SI/SLEA), and BST bilaterally [67]. Consistent with recent recommendations [68,69], the ROI was created using a combination of the Mai and Harvard-Oxford atlases [70-74] and included the probabilistic BST ROI developed by Theiss and colleagues ($p$>0%) [75] and the Harvard-Oxford probabilistic amygdala ($p$>50%). Using this as a starting point, voxels in the region of the SI/SLEA was manually added in the coronal plane of the 1-mm MNI152 template, working from rostral to caudal, and confirmed in the other planes. At intermediate levels of the amygdala's rostral-caudal axis, where the BST was no longer visible, the SI/SLEA was limited to voxels dorsal to the amygdala and ventral to the putamen and pallidum. SI/SLEA voxels were included until the head of the hippocampus was clearly visible. Voxels in neighboring regions of the accumbens, caudate, putamen, pallidum, thalamus, and ventricles (Harvard-Oxford atlas, $p$>50%) were excluded using a Boolean 'NOT.' The resulting bilateral ROI was decimated to 2-mm$^3$ (total: 1,205 voxels; 9,640 mm$^3$). For illustrative purposes, the 1-mm ROI is shown.

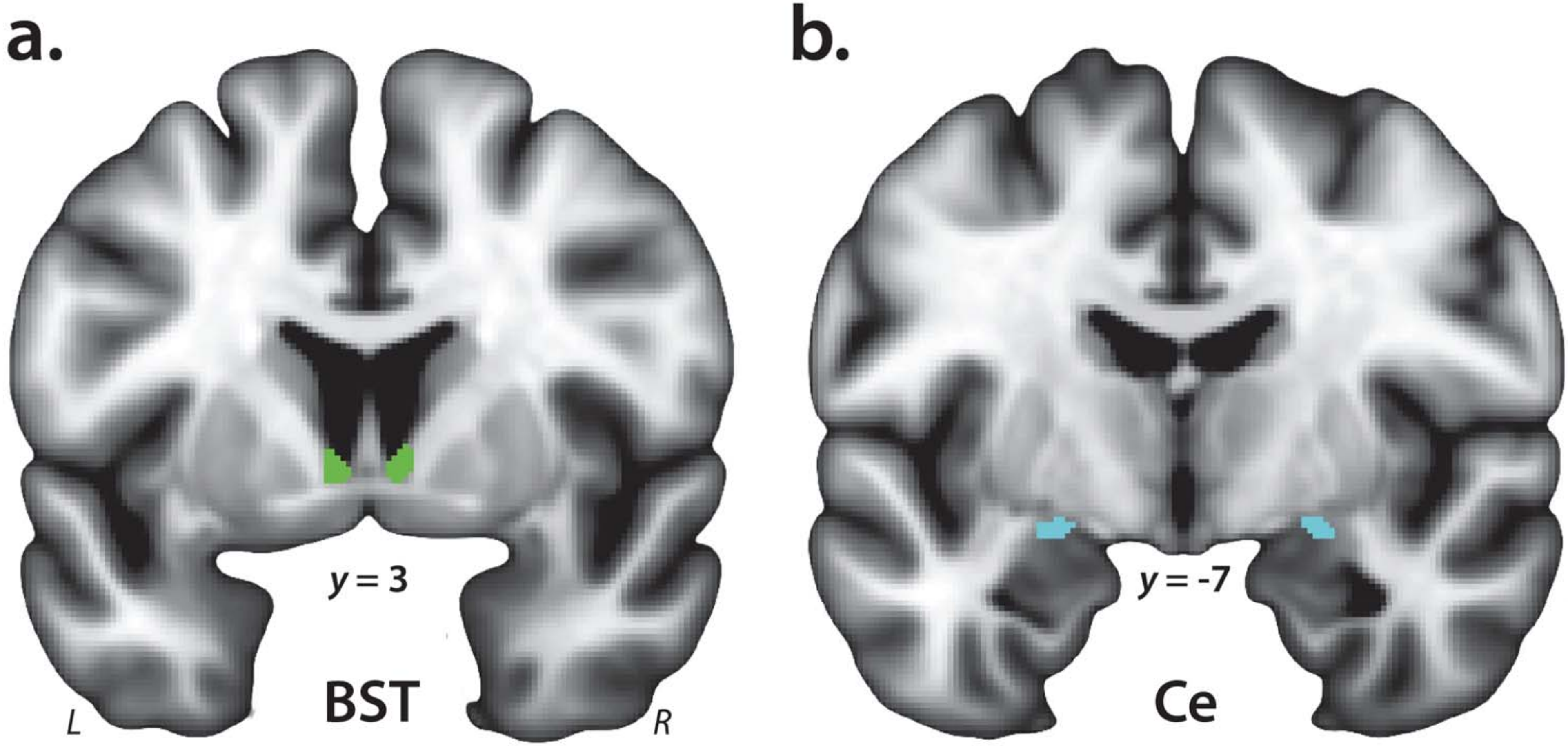

**Supplementary Figure S3. BST and Ce ROIs.** ***a. BST.*** The derivation of the probabilistic BST ROI (*green*) is detailed in [81] and was thresholded at 25%. The seed mostly encompasses the supra-commissural BST, given the difficulty of reliably discriminating the borders of regions below the anterior commissure on the basis of T1-weighted images [cf. 82]. ***b. Ce.*** The derivation of the Ce ROI (*cyan*) is described in more detail in Tillman et al. (2017). For illustrative purposes, 1-mm ROIs are shown. Analyses employed ROIs decimated to the 2-mm resolution of the EPI data. Single-subject data were visually inspected to ensure that the ROIs were correctly aligned to the spatially normalized T1 images. Abbreviations—BST, bed nucleus of the stria terminalis; Ce, central nucleus of the amygdala; L, left hemisphere; R, right hemisphere.

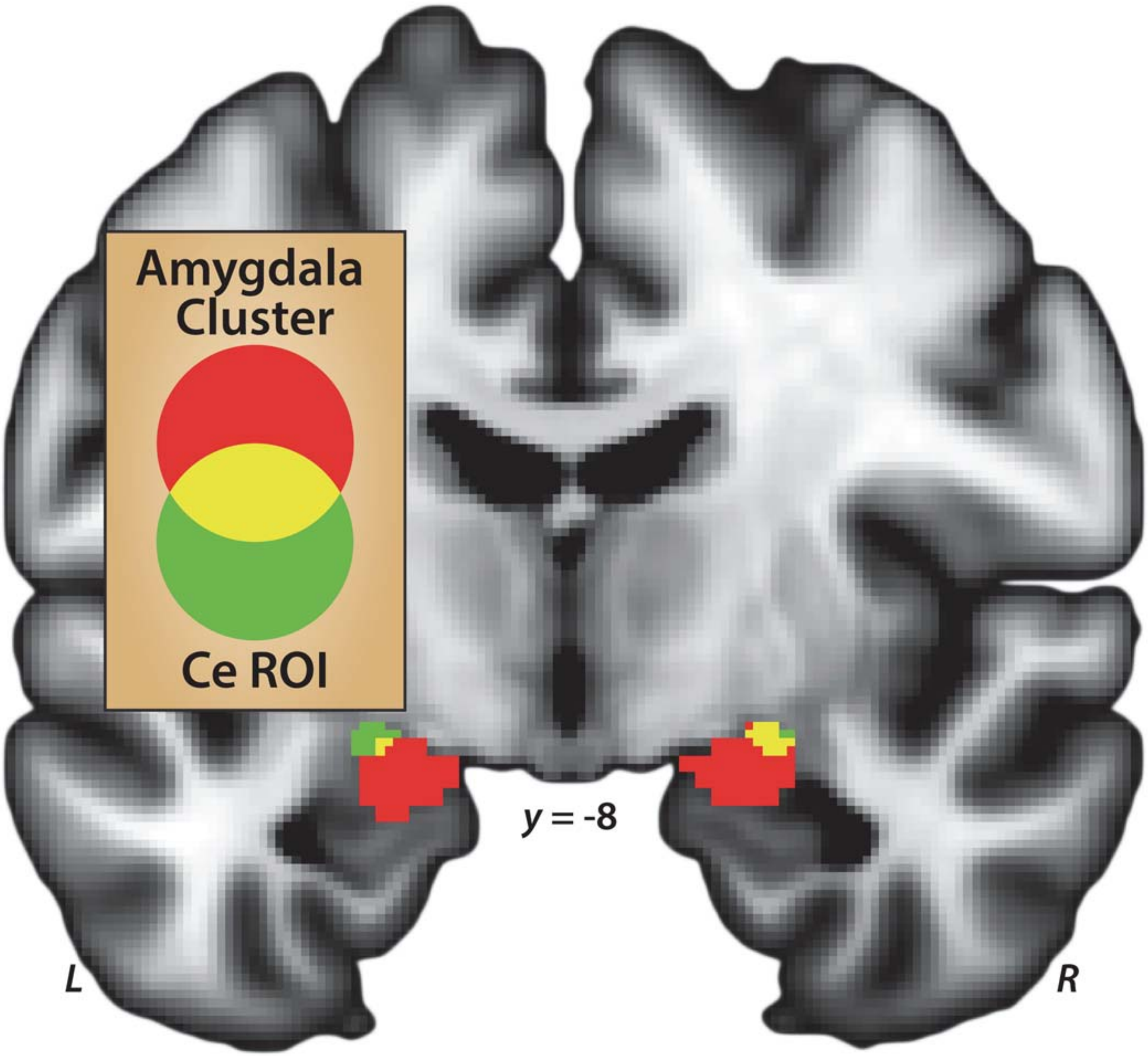

**Supplementary Figure S4.** ***The amygdala cluster identified in voxelwise analyses overlaps the anatomically defined Ce ROI.*** While the cluster included numerous amygdala nuclei, the left (*x*=-20, *y*=-10, *z*=-14) and right (*x*=22, *y*=-8, *z*=-16) peaks lie in the dorsocaudal region where the more dorsal Ce and more ventral basomedial nuclei abut. The derivation of the Ce seed (*cyan*) is described in more detail in Tillman et al. (2018). Abbreviations—Ce, central nucleus of the amygdala; L, left hemisphere; R, right hemisphere; ROI, region of interest.

**Supplementary Table 1***. Descriptive statistics for clusters identified by the faces vs. places contrast using p<.05, small-volume corrected*[a]

| | | mm³ | *t* | *x* | *y* | *z* |
|---|---|---|---|---|---|---|
| | ***Faces > Places*** | | | | | |
| | R Frontal | 13,824 | | | | |
| | R Frontal Pole[b] | | 4.67 | 50 | 38 | 28 |
| | R Inferior Frontal Gyrus, pars opercularis[b] | | 8.92 | 54 | 12 | 22 |
| | R Precentral Gyrus[b] | | 6.53 | 44 | -2 | 40 |
| | L Insular Cortex[b] | 696 | 6.15 | -34 | 24 | 0 |
| | R Frontal Operculum Cortex[b] | 768 | 6.65 | 38 | 24 | 0 |
| | R Superior Frontal Gyrus[b] | 344 | 5.05 | 6 | 18 | 56 |
| | L Inferior Frontal Gyrus[b] | 328 | 4.93 | -44 | 16 | 26 |
| | L Temporal/Amygdala | 2,480 | | | | |
| | L Temporal Pole[b] | | 5.48 | -34 | 4 | -28 |
| | L Parahippocampal Gyrus, anterior[b] | | 6.05 | -32 | 0 | -32 |
| | L Dorsal Amygdala[c] | | 12.59 | -20 | -10 | -14 |
| | R Posterior Temporal/Amygdala | 2,832 | | | | |
| | R Temporal Fusiform Cortex, anterior[b] | | 5.41 | 32 | -2 | -34 |
| | R Inferior Temporal Gyrus, anterior[b] | | 5.94 | 40 | -2 | -40 |

| | | | | | | |
|---|---|---|---|---|---|---|
| | R Dorsal Amygdala[c] | | 12.22 | 22 | -8 | -16 |
| | R Thalamus[b] | 88 | 4.64 | 6 | -4 | 0 |
| | L Postcentral Gyrus[b] | 1,208 | 7.39 | -48 | -18 | 48 |
| | R Temporal-Occipital | 14,696 | | | | |
| | R Middle Temporal Gyrus, posterior[b] | | 6.32 | 50 | -26 | -4 |
| | R Middle Temporal Gyrus, temporooccipital part[b] | | 10.29 | 54 | -60 | 10 |
| | R Lateral Occipital Cortex, inferior[b] | | 11.51 | 54 | -68 | 6 |
| | R Supramarginal Gyrus, anterior[b] | 352 | 4.78 | 54 | -34 | 52 |
| | R Occipital-Temporal | 3,184 | | | | |
| | R Inferior Temporal Gyrus[b] | | 7.75 | 46 | -40 | -18 |
| | R Temporal Occipital Fusiform Cortex[b] | | 9.81 | 42 | -48 | -20 |
| | L Temporal Occipital Fusiform Cortex[b] | 2,416 | 10.22 | -40 | -50 | -18 |
| | L Occipital-Temporal | 5,040 | | | | |
| | L Middle Temporal Gyrus, temporooccipital part[b] | | 6.08 | -58 | -52 | 10 |
| | L Lateral Occipital Cortex, inferior[b] | | 7.19 | -52 | -68 | 8 |
| | L Intracalcarine Cortex[b] | 128 | 4.67 | -8 | -88 | 10 |
| | ***Places > Faces*** | | | | | |
| | Bilateral Frontal | 6,808 | | | | |

| | | | | | | |
|---|---|---|---|---|---|---|
| | L Frontal Pole[b] | | 4.85 | -2 | 62 | 2 |
| | R Frontal Pole[b] | | 4.78 | 6 | 58 | 2 |
| | L Cingulate Gyrus, anterior[b] | | 6.03 | -4 | 40 | -4 |
| | R Paracingulate Gyrus[b] | | 6.05 | 6 | 40 | -6 |
| | R Cingulate Gyrus, anterior[b] | | 5.83 | 4 | 38 | 18 |
| | L Paracingulate Gyrus[b] | | 5.58 | -10 | 38 | -6 |
| | L Frontal Pole[b] | 120 | 5.01 | -22 | 50 | 28 |
| | L Frontal | 2,816 | | | | |
| | L Frontal Pole[b] | | 5.38 | -18 | 40 | 40 |
| | L Middle Frontal Gyrus[b] | | 4.67 | -28 | 26 | 40 |
| | L Superior Frontal Gyrus[b] | | 6.69 | -18 | 26 | 48 |
| | L Frontal Pole[b] | 384 | 5.49 | -34 | 38 | -10 |
| | R Frontal Pole[b] | 128 | 4.67 | 32 | 38 | -10 |
| | R Middle Frontal Gyrus[b] | 256 | 4.87 | 28 | 26 | 36 |
| | R Cingulate Gyrus, anterior[b] | 248 | 4.47 | 2 | 6 | 32 |
| | L Insular Cortex[b] | 304 | 5.34 | -44 | 4 | -6 |
| | R Temporal-Parietal | 4,248 | | | | |
| | R Precentral Gyrus[b] | | 4.71 | 62 | 0 | 10 |

| | | | | | | |
|---|---|---|---|---|---|---|
| | R Superior Temporal Gyrus, posterior[b] | | 5.78 | 66 | -20 | 4 |
| | R Planum Temporale[b] | | 6.14 | 64 | -26 | 12 |
| | R Parietal Operculum Cortex[b] | | 6.58 | 50 | -28 | 24 |
| | L Temporal | 880 | | | | |
| | L Middle Temporal Gyrus, anterior[b] | | 5.31 | -62 | -2 | -12 |
| | L Superior Temporal Gyrus, anterior[b] | | 4.45 | -60 | -6 | -6 |
| | L Middle Temporal Gyrus, posterior[b] | | 5.23 | -66 | -18 | -10 |
| | L Temporal | 728 | | | | |
| | L Superior Temporal Gyrus, anterior[b] | | 5.40 | -62 | -4 | 4 |
| | L Central Opercular Cortex[b] | | 4.59 | -56 | -6 | 10 |
| | R Inferior Visual Cortex | 188,488 | | | | |
| | R Parahippocampal Gyrus, posterior[b] | | 10.09 | 20 | -24 | -20 |
| | R Thalamus[b] | | 8.31 | 24 | -32 | 4 |
| | L Thalamus[b] | | 9.55 | -20 | -32 | -2 |
| | L Parahippocampal Gyrus, anterior[b] | | 15.72 | -24 | -38 | -14 |
| | R Lingual Gyrus[b] | | 20.56 | 24 | -40 | -12 |
| | R Cingulate Gyrus, posterior[b] | | 11.06 | 10 | -46 | 4 |
| | L Temporal Occipital Fusiform Cortex[b] | | 19.16 | -28 | -50 | -10 |

| | | | | | | |
|---|---|---|---|---|---|---|
| | L Lingual Gyrus[b] | | 12.50 | -16 | -52 | 2 |
| | R Temporal Occipital Fusiform Cortex[b] | | 22.04 | 30 | -54 | -10 |
| | R Precuneus Cortex[b] | | 16.79 | 18 | -58 | 12 |
| | L Precuneus Cortex[b] | | 16.27 | -16 | -60 | 8 |
| | L Occipital Fusiform Gyrus[b] | | 17.06 | -26 | -70 | -12 |
| | R Occipital Fusiform Gyrus[b] | | 17.21 | 26 | -72 | -8 |
| | L Lateral Occipital Cortex, superior [b] | | 9.52 | -22 | -84 | 38 |
| | R Lateral Occipital Cortex, superior[b] | | 14.29 | 40 | -86 | 12 |
| | L Occipital Pole[b] | | 14.11 | -30 | -92 | 16 |
| | R Occipital Pole[b] | | 13.53 | 18 | -98 | 0 |
| | L Parietal Operculum Cortex[b] | 336 | 4.96 | -60 | -28 | 16 |
| | L Angular Gyrus[b] | 384 | 5.38 | -50 | -56 | 30 |
| | L Middle Temporal Gyrus, temporooccipital[b] | 1,912 | 6.94 | -58 | -58 | -8 |

[a] The faces-vs.-places contrast was thresholded at $p<.05$ FWE-corrected for the extent of the EAc ROI (1,205 voxels; 9,640 $mm^3$). For transparency, all clusters greater than or equal to 80 $mm^3$ (10 native EPI voxels) are reported. [b] Lies outside the *a priori* EAc ROI and is not significant. [c] Lies inside the *a priori* EAc ROI and is significant.

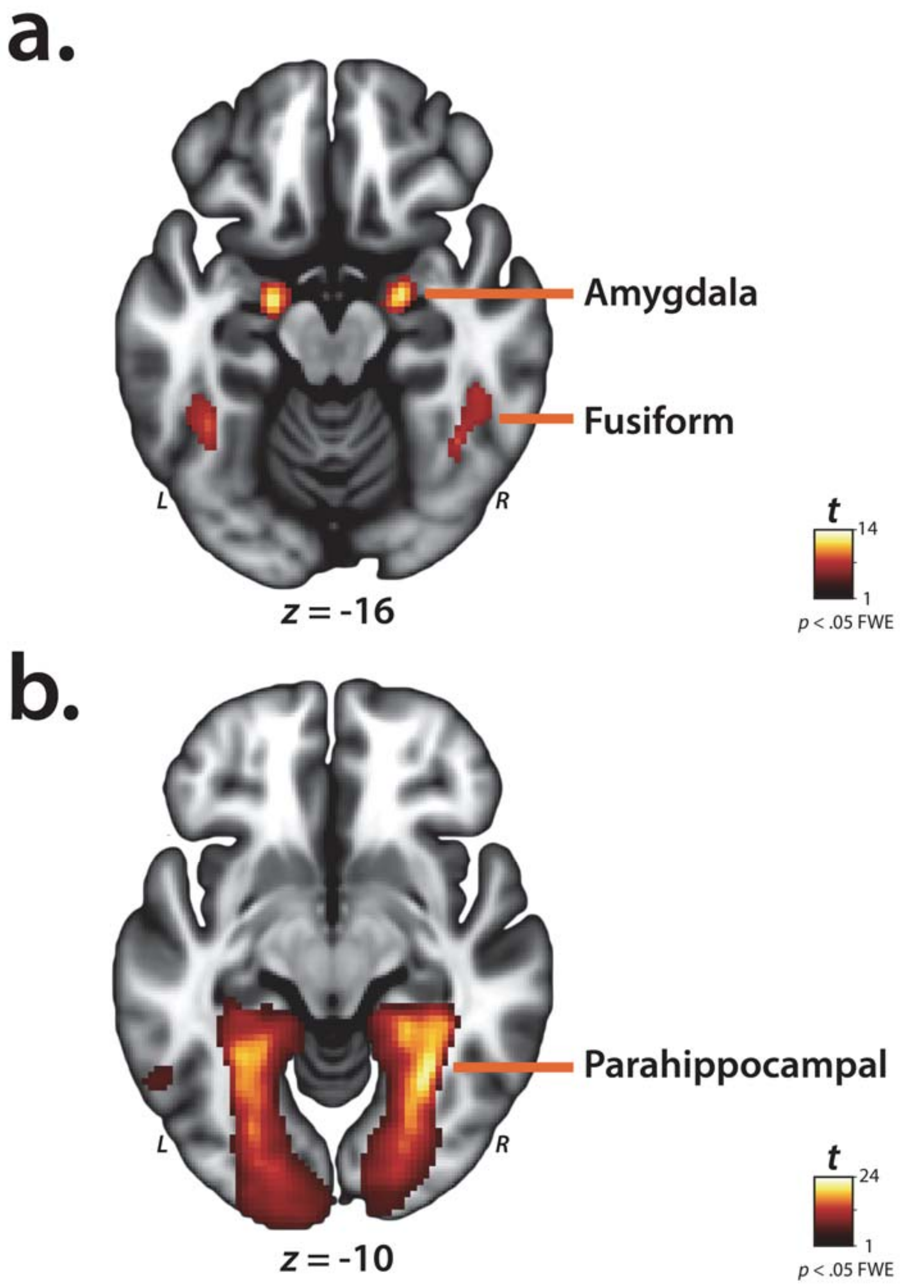

**Supplementary Figure S5.** ***Regions identified by a whole-brain, voxelwise regression analysis (p < .05, whole-brain FWE corrected). A.*** The amygdala and fusiform cortex show significantly greater activation to threat-related faces than buildings***. B.*** The parahippocampal cortex shows significantly greater activation to buildings than threat-related faces***.*** Abbreviations—L, left hemisphere; R, right hemisphere.

**Supplementary Table 2.** ***Descriptive statistics for clusters identified by the faces vs. places contrast using p<.05, whole-brain corrected[a]***

| | | mm³ | *t* | *x* | *y* | *z* |
|---|---|---|---|---|---|---|
| | ***Faces > Houses*** | | | | | |
| | R Frontal Operculum Cortex | 80 | 6.65 | 38 | 24 | 0 |
| | R Inferior Frontal Gyrus, pars opercularis | 3,640 | 8.92 | 54 | 12 | 22 |
| | R Precentral Gyrus | 224 | 6.53 | 44 | -2 | 40 |
| | L Amygdala | 800 | 12.59 | -20 | -10 | -14 |
| | R Amygdala | 920 | 12.22 | 22 | -8 | -16 |
| | L Precentral | 360 | | | | |
| | L Postcentral Gyrus | | 7.39 | -48 | -18 | 48 |
| | L Precentral Gyrus | | 6.09 | -36 | -22 | 50 |
| | R Temporal-Occipital | 5,936 | | | | |
| | R Middle Temporal Gyrus, posterior | | 6.32 | 50 | -26 | -4 |
| | R Middle Temporal Gyrus, temporooccipital part | | 7.69 | 54 | -40 | 2 |
| | R Lateral Occipital Cortex, inferior | | 11.51 | 54 | -68 | 6 |
| | R Temporal-Occipital | 1,536 | | | | |
| | R Inferior Temporal Gyrus, temporooccipital part | | 7.75 | 46 | -40 | -18 |

| | | | | | | |
|---|---|---|---|---|---|---|
| | R Temporal Occipital Fusiform Cortex | | 9.81 | 42 | -48 | -20 |
| | L Temporal Occipital Fusiform Cortex | 1,144 | 10.22 | -40 | -50 | -18 |
| | L Temporal-Occipital | 1,000 | | | | |
| | L Middle Temporal Gyrus, temporooccipital part | | 6.08 | -58 | -52 | 10 |
| | L Lateral Occipital Cortex, inferior | | 7.19 | -52 | -68 | 8 |
| | ***Places > Faces*** | | | | | |
| | L Superior Frontal Gyrus (C7) | 184 | 6.69 | -18 | 26 | 48 |
| | R Inferior Visual Cortex | 55,360 | | | | |
| | R Hippocampus | | 8.77 | 30 | -20 | -22 |
| | R Parahippocampal Gyrus | | 10.09 | 20 | -24 | -20 |
| | R Lingual Gyrus | | 20.56 | 24 | -40 | -12 |
| | R Temporal Occipital Fusiform Cortex | | 22.04 | 30 | -54 | -10 |
| | R Lateral Occipital Cortex, superior | | 15.54 | 36 | -88 | 16 |
| | L Thalamus | 288 | 9.55 | -20 | -32 | -2 |
| | R Thalamus | 240 | 8.31 | 24 | -32 | 4 |
| | L Posterior Cingulate/Precuneus | 4,704 | | | | |
| | L Cingulate Gyrus, posterior | | 8.38 | -6 | -34 | 36 |
| | L Precuneus Cortex | | 7.97 | -8 | -42 | 44 |
| | L Inferior Visual Cortex | 60,288 | | | | |

| | | | | | | |
|---|---|---|---|---|---|---|
| | L Parahippocampal Gyrus, posterior | | 15.72 | -24 | -38 | -14 |
| | L Temporal Occipital Fusiform Gyrus | | 19.49 | -28 | -52 | -6 |
| | L Cingulate Gyrus, posterior | | 12.50 | -16 | -52 | 2 |
| | L Precuneous Cortex | | 16.27 | -16 | -60 | 8 |
| | L Occipital Fusiform Gyrus | | 17.06 | -26 | -70 | -12 |
| | L Lateral Occipital Cortex, superior | | 9.52 | -22 | -84 | 38 |
| | L Occipital Pole | | 14.11 | -30 | -92 | 16 |
| | L Middle Temporal Gyrus | 424 | 6.94 | -58 | -58 | -8 |

[a]Clusters greater than or equal to 80 $mm^3$ (10 native EPI voxels) are reported.

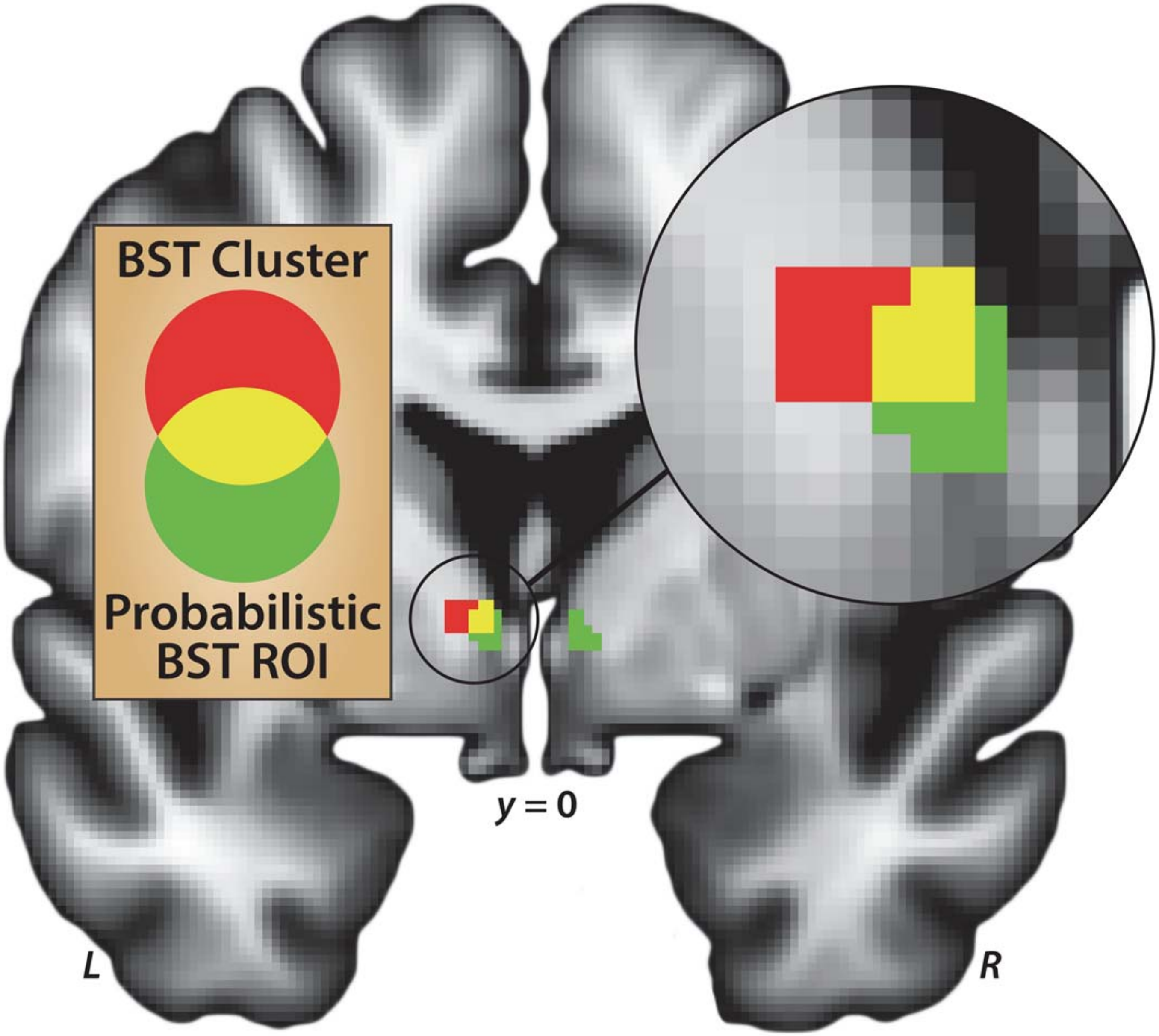

**Supplementary Figure S6.** ***The BST cluster identified in voxelwise analyses overlaps the anatomically defined BST ROI.*** The derivation of the probabilistic BST ROI (*green*) is detailed in [81]. The same pattern was evident using other available BST ROIs (not shown) [83,84]. Abbreviations—L, left hemisphere; R, right hemisphere; ROI, region of interest.

**Supplementary Table 3.** ***Descriptive statistics for clusters identified by the Group × Condition contrast using p<.05, small-volume corrected*** [a]

| | | mm³ | *t* | *x* | *y* | *z* |
|---|---|---|---|---|---|---|
| | ***Alcohol < Placebo: Faces minus Places*** | | | | | |
| | L Insula | 168 | | | | |
| | L Frontal Operculum Cortex[b] | | 4.19 | -32 | 22 | 8 |
| | L Insular Cortex[b] | | 5.46 | -34 | 20 | 2 |
| | L BST[c] | 104 | 5.46 | -8 | -2 | 0 |
| | L Thalamus[b] | 80 | 5.15 | -6 | -24 | 0 |
| | ***Alcohol > Placebo: Faces minus Places*** | | | | | |
| | R Temporal Occipital Fusiform Cortex[b] | 360 | 5.94 | 26 | -58 | -12 |
| | R Lateral Occipital Cortex, superior[b] | 200 | 5.10 | 32 | -80 | 12 |

[a]The Group × Condition interaction contrast was thresholded at *p*<.05 FWE-corrected for the extent of the EAc ROI (1,205 voxels; 9,640 mm$^3$). For transparency, all clusters greater than or equal to 80 mm$^3$ (10 native EPI voxels) are reported. [b]Lies outside the *a priori* EAc ROI and is not significant. [c]Lies inside the *a priori* EAc ROI and is significant.

### Analyses Controlling for Group Differences in Performance

On average, subjects were highly accurate at performing the simple discrimination tasks (**Supplementary Table 4**). Nevertheless, performance was ~8% lower in the alcohol compared to the placebo group.

**Supplementary Table 4:** ***Mean Percentage Correct (SD) for the Complete Sample***

| | **Total (*N*=49)** | **Placebo (*N*=22)** | **Alcohol (*N*=27)** |
|---|---|---|---|
| Faces | 84.4 (8.4) | 89.1 (5.3) | 80.6 (8.5) |
| Places | 88.8 (8.8) | 93.0 (5.0) | 85.5 (9.8) |
| Overall | 86.8 (7.9) | 91.1 (4.9) | 83.2 (8.2) |

To ascertain whether key neural effects were primarily driven by group differences in task engagement, imaging data were re-analyzed using a subset of subjects selected to have matching levels of overall performance on the face/place discrimination task (*n* = 15/group). The least-accurate subjects from the alcohol group and the most-accurate subjects from the placebo group were iteratively excluded until the mean of the alcohol group numerically exceeded that of the placebo group, as detailed in **Supplementary Table 5**.

**Supplementary Table 5:** ***Mean Percentage Correct (SD) for the Performance-Matched Sub-Sample***

| | **Total (*N*=30)** | **Placebo (*N*=15)** | **Alcohol (*N*=15)** |
|---|---|---|---|
| Faces | 86.9 (0.05) | 86.9 (0.05) | 86.9 (0.05) |
| Places | 91.1 (0.04) | 91.0 (0.05) | 91.2 (0.03) |
| Total Accuracy | 89.0 (0.04) | 88.9 (0.05) | 89.1 (0.04) |

Using this sub-sample ($N$=30), acute alcohol administration was still associated with a substantial reduction in threat-related activation in the left BST/thalamus ($t(29)=3.51$, $p=.001$, uncorrected; $x=-8$, $y=-2$, $z=-2$). A similar effect was observed at the location of the original BST peak ($t(29)=3.35$, $p=.002$, uncorrected; $x=-8$, $y=-2$, $z=0$).

With regard to the anatomically defined, *a priori* ROI analyses, in the intact sample ($n$=49), a mixed-model GLM revealed greater threat-related activation in the Ce relative to the BST ($F(1,47)=32.99$, $p<.001$) and a near-significant alcohol-dampening effect across regions ($F_{W-S}(1,47)=3.93$, $p=.053$). Other omnibus effects were not significant ($ps>.15$). Likewise, in the performance-matched sub-sample ($N$=30), results again revealed greater threat-related activation in the Ce relative to the BST ($F(1,28)=30.58$, $p<.001$) and a trend-level alcohol-dampening effect across regions ($F_{W-S}(1,47)=3.17$, $p=.088$). The size of the threat-dampening effect was numerically larger in the sub-sample ($p\eta^2=.102$) compared to the full sample ($p\eta^2=.075$). Collectively, these results suggest that our imaging results are not primarily driven by group differences in overt behavioral performance, a proxy for overall task engagement.
